# Conjugative IncC plasmid entry triggers the SOS response and promotes effective transfer of the integrative antibiotic resistance element SGI1

**DOI:** 10.1101/2022.06.09.495587

**Authors:** Marine C. Pons, Karine Praud, Sandra Da Re, Axel Cloeckaert, Benoît Doublet

## Abstract

The broad host range IncC plasmid family and the integrative mobilizable *Salmonella* Genomic Island 1 (SGI1) and its derivatives enable the spread of medically-important antibiotic resistance genes among Gram-negative pathogens. Although several aspects of the complex functional interactions between IncC plasmids and SGI1 have been recently deciphered regarding their conjugative transfer and incompatibility, the biological signal resulting in the hijacking of the conjugative plasmid by the integrative mobilizable element remains unknown. Here, we demonstrate that the conjugative entry of IncC/IncA plasmids is detected at an early stage by SGI1 through the transient activation of the SOS response, which induces the expression of the SGI1 master activators SgaDC, shown to play a crucial role in the complex biology between SGI1 and IncC plasmids. Besides, we developed an original tripartite conjugation approach to directly monitor SGI1 mobilization in a time-dependent manner following conjugative entry of IncC plasmids. Finally, we propose an updated biological model of the conjugative mobilization of the chromosomal resistance element SGI1 by IncC plasmids.

**IMPORTANCE:** Antimicrobial resistance has become a major public health issue, particularly with the increase in multidrug resistance (MDR) in both animal and human pathogenic bacteria, and with the emergence of resistance to medically important antibiotics. The spread between bacteria of successful mobile genetic elements such as conjugative plasmids and integrative elements conferring multidrug resistance is the main driving force in the dissemination of acquired antibiotic resistances among Gram-negative bacteria. Broad-host range IncC plasmids and their integrative mobilizable SGI1 counterparts contribute to the spread of critically-important resistance genes (e.g., ESBLs, and carbapenemases). A better knowledge of the complex biology of these broad-host range mobile elements will help to understand the dissemination of antimicrobial resistance genes that occurred across *γ-proteobacteria* borders.

## INTRODUCTION

Mobile genetic elements play an essential role in the emergence and dissemination of antibiotic resistance among bacteria (1, 2). Among them, self-transmissible IncC plasmids are broad-host-range conjugative plasmids ranging from 100 to 200 kb that contribute to the dissemination of numerous antibiotic resistance genes in the *γ-proteobacteria* (1–3). Moreover, IncC plasmids have been shown to serve as helper conjugative plasmids to drive transfer of diverse mobilizable genomic islands (also named integrative mobilizable element) that are integrated into the chromosome and that can carry also medically-important antibiotic resistance genes (3, 5–12). SGI1 is the prototype of a large family of multidrug resistance integrative mobilizable elements that are conjugally mobilized *in trans* by plasmids of the IncC/IncA family (7, 13, 14). SGI1 and IncC plasmids share a complex biology regarding their conjugative transfer and incompatibility (15–23). SGI1 encodes a few transfer proteins that reshape the IncC-encoded mating pore in order to promote its own dissemination in bacterial populations already harbouring an IncC plasmid (16). Besides, the SGI1 Toxin-Antitoxin system SgiAT and replication of excised SGI1 have been shown to participate in SGI1 stability and in the concomitant destabilization of the IncC plasmids (17, 19, 20). Moreover, IncC plasmids and SGI1 share a complex transcriptional regulatory network, each element having homologous master activators, i.e. AcaDC and SgaDC, respectively, which have been shown to activate the same regulons in both elements (15, 18, 19, 21). Durand et al. recently demonstrated that SgaDC and AcaDC activate the expression of AcaB, an IncC transcriptional regulator which in turn in cooperation with AcaDC have been shown to trigger the expression of the conjugative IncC machinery through a positive feedback loop of mutual expression (18, 24).

Conjugative circular plasmids are considered to horizontally transfer to plasmidless recipient cells mainly as single-stranded DNA (ssDNA) (25). The transient occurrence of ssDNA in the recipient cell during bacterial conjugation generally leads to the induction of the SOS response (26). Numerous physical or chemical treatments (UV irradiation, antibiotics, i.e. ciprofloxacin, trimethoprim, or DNA cross-linking agent such as mitomycin C) can trigger the accumulation of ssDNA and subsequently the activation of the bacterial SOS response (26–28). Beyond the host-encoded SOS-regulon mainly responding to DNA damage, several genes carried by mobile genetic elements, such as phages and integrative elements, are also regulated by the SOS response, either directly by the host-encoded LexA repressor or by their own analogous repressors (28, 29). These SOS-regulated genes are involved in horizontal gene transfer (e.g., antibiotic resistance), bacterial virulence, and evolution (28). To the best of our knowledge, plasmids of the IncC/IncA family have not yet been reported to conjugally transfer as ssDNA, nor to activate the SOS response following conjugative entry in recipient cells (26, 30).

## RESULTS

### The SOS response is induced by conjugative entry of IncC plasmids in recipient cells

In order to test whether the conjugative transfer of IncC/IncA plasmids induces the SOS response, we performed a SOS β-galactosidase reporter assay during conjugation with the reference conjugative plasmids IncC-R55, IncA-RA1, and IncW-Rsa (positive control) in *Escherichia coli* used as recipient cells (26). SOS induction in the recipient *E. coli* population was measured using the P*recN*::*lacZ* transcriptional fusion as reporter in conjugation assays. Knowing that plasmids have different kinetics of transfer frequency, the conjugation frequencies were determined at different time points after donor and recipient contact to identify the earliest time point allowing transfer frequencies of at least ∼10^-4^-10^-3^ (arbitrary choice; Fig. 1A). We then studied the SOS induction through β-galactosidase activity in the recipient population at this earliest time, i.e. at 1h for IncC-R55 and IncA-RA1 plasmids and 2h for IncW-Rsa plasmid (Fig. 1A). The basal β-galactosidase activity was determined per recipient cell in absence of conjugative plasmid (empty donor, no SOS induction by ssDNA entry) (Fig. 1B) and the specific β-galactosidase activity per transconjugant for each plasmid (Fig. 1C). SOS induction (expressed as the induction ratio between specific β-galactosidase activity per transconjugant cell and basal β-galactosidase activity per recipient cell; see Methods) was observed in all mating assays with 8050-fold, 2412-fold and 183-fold induction ratio for IncW-Rsa, IncC-R55 and IncA-RA1 plasmids, respectively (Fig. 1D). The difference of SOS induction between these plasmids can be explained by asynchronous transfers, and different transfer rates (Fig. 1A), resulting in asynchronous SOS induction in the transconjugant population. Using a *recA^-^* recipient strain in the same conjugation assay, β-galactosidase activity in the recipient population remained at basal levels strongly suggesting that ssDNA entry of plasmids is responsible of the SOS induction through activation of LexA auto-proteolysis by RecA-ssDNA complex. Thus, as previously described for other plasmid families, these results confirmed that the transfer of IncC/IncA plasmid families induces transiently the SOS response in the recipient cell following conjugative entry (26).

**FIG 1.**
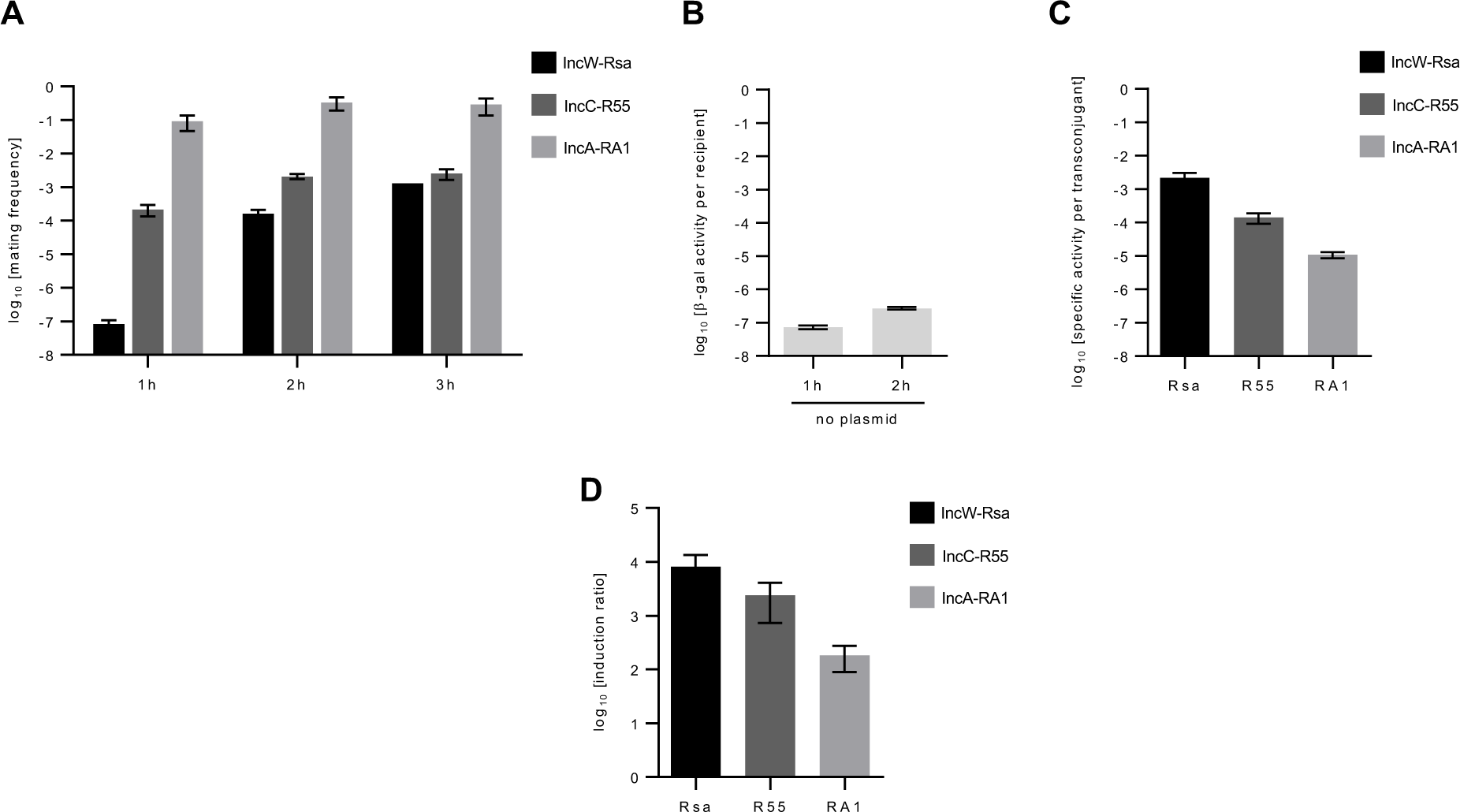
Conjugative entry of IncC/IncA plasmids activates the SOS response in the recipient cells. The specific activation of the SOS response following plasmid entry in recipient cells was determined in conjugation assays using donor *E. coli* strains TOP10 harbouring conjugative plasmids and recipient *E. coli* strain MG1655 carrying reporter vectors (*lacZ* expression under the promoter P*recN,* pMP002 or pMP010; see details in methods section) (**A**), Time-dependent transfer frequency of Rsa (IncW, positive control), R55 (IncC) and RA1 (IncA) plasmids. Transfer frequencies are expressed as the number of transconjugant per donor CFUs. (**B**), basal β-galactosidase activity per recipient without conjugative plasmid in the donor. (**C**), Specific β-galactosidase activity per transconjugant for each plasmid at the time point corresponding to a transfer frequency of at least 10^-4^ (R55 and RA1: 1h; Rsa: 2h). (**D**), SOS induction ratio correspond to the specific β-galactosidase activity per transconjugant divided by the basal β-galactosidase activity in recipient. In panels (**B**), (**C**) and (**D**), β-galactosidase activities and induction ratio were calculated as described in Materials and Methods. The bars represent the mean with standard error of the mean obtained from three independent experiments.

### Expression of the SGI1 master activators *sgaDC* is activated by the SOS response

Using the regulatory sequence analysis tools (http://embnet.ccg.unam.mx/rsat/matrix-scan_form.cgi) with the RegulonDB database, we identified 6 putative LexA binding sites in the SGI1 backbone sequence of SGI1-C from *S.* Agona strain 47SA97 (considered as reference, and hereafter named SGI1 and used in all experiments), among which 2 putative sites were located in promoter regions of the master activator genes *sgaDC* and the toxin-antitoxin genes *sgiAT* (Table 1, Fig. 2) (31, 32). We therefore hypothesized that the induction of the SOS response could be detected by SGI1 as a signal for horizontal transfer, just after the conjugative entry of IncC plasmids in SGI1 bearing cells.

**FIG 2.**
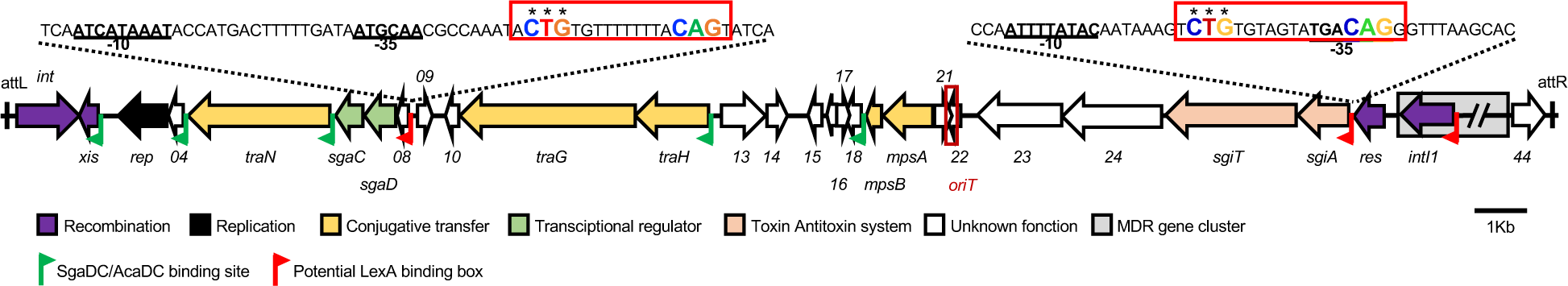
Linear schematic representation of conserved SGI1 backbone. Integrated SGI1 is flanked by *attL* and *attR* attachment sites. The position and orientation of open reading frames (ORFs) are indicated by arrows. ORF functions from predictions or previous functional analyses are color-coded. Putative LexA binding boxes are represented as red flags. Partial sequences of *sgaDC* and *sgiAT* promoters are indicated showing putative -10, -35 regions, and LexA binding box (red box). Stars indicate the 3 essential nucleotides (CTG) of LexA binding boxes that have been substituted in the mutated probes for electrophoretic mobility shift assay (Figs. 3 & S2). SgaDC/AcaDC binding sites are indicated by green flags. *oriT* represents the SGI1 origin of transfer.

**TABLE 1.**
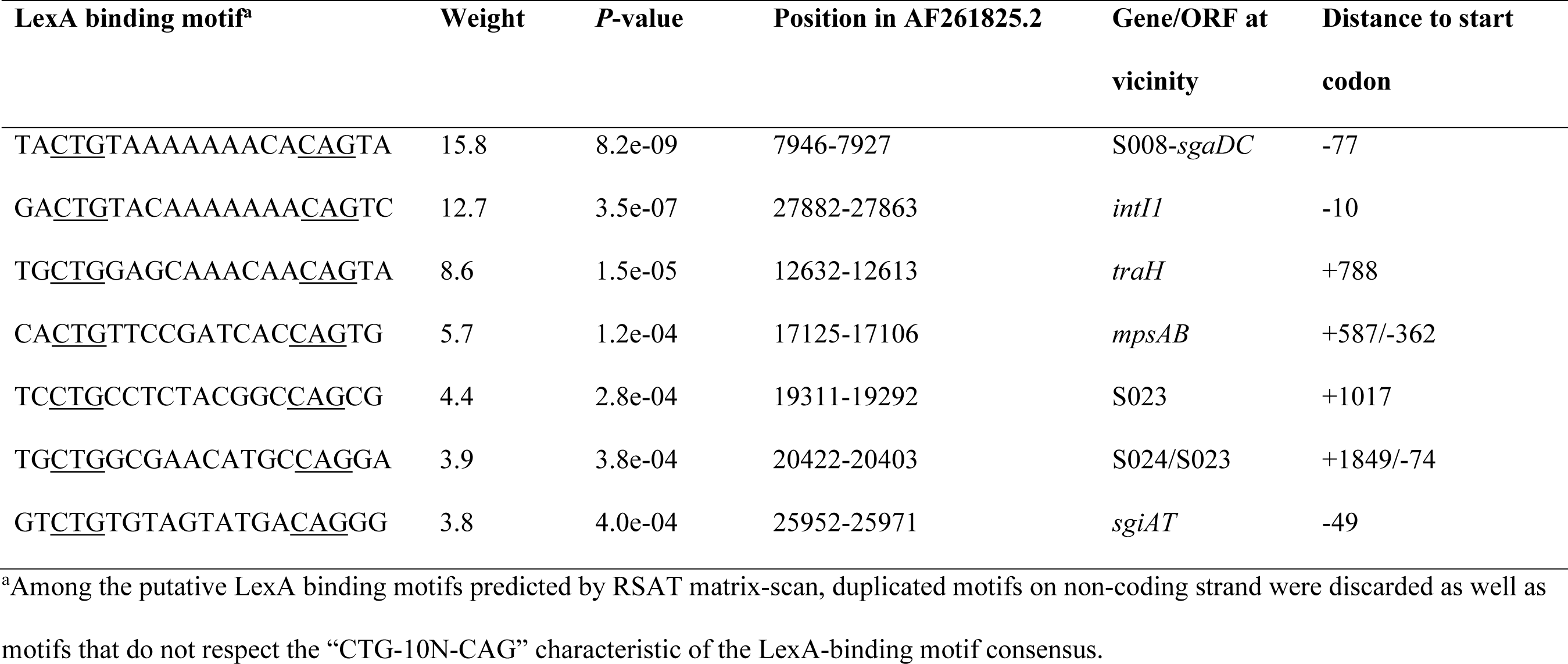
Putative LexA binding si tes in SGI1 identified by the regulatory sequence analysis tools (http://embnet.ccg.unam.mx/rsat/matrix-scan_form.cgi) using the RegulonDB database mong the putative LexA binding motifs predicted by RSAT matrix-scan, duplicated motifs on non-coding strand were discarded as well as tifs that do not respect the “CTG-10N-CAG” characteristic of the LexA-binding motif consensus.

Assessment of the P*sgaDC* and P*sgiAT* promoter activities under SOS inducing conditions (presence of ciprofloxacin or mitomycin C) was performed using *lacZ* transcriptional fusions with these promoters in two *Salmonella enterica* serotypes, Agona and Kentucky ST198, and in *E. coli* through β-galactosidase assays. P*recN* and P*acaDC* were used as positive and negative controls, respectively (Fig. S1) (8, 32). The activity of P*sgaDC* was low in the absence of treatment and induced in the presence of ciprofloxacin and mitomycin C. It resulted respectively in 5.5-fold and 9.2-fold increase of P*sgaDC* activity in *S.* Kentucky ST198, and in 6.1-fold increase for both stress in *S.* Agona (Fig. 3A). A much lower but still significant, increase of P*sgaDC* activity was also observed in *E. coli* background after ciprofloxacin and mitomycin C treatments (Fig. 3B). To confirm these results suggesting that P*sgaDC* activity is induced by the SOS response, we performed the same β-galactosidase reporter assays in *E. coli* mutants for the SOS response (SOS^OFF^: *lexA3* mutant coding for a non-cleavable LexA protein, and SOS^ON^: *lexA51* mutant coding for LexA protein variant unable to bind on its binding box). P*sgaDC* activity showed a 4.7-fold increase in the *E. coli* SOS^ON^ mutant compared to the wild-type *E. coli* strain MG1655 without chemical treatment (Fig. 3B). Similar results were obtained for P*sgaDC* β-galactosidase activity in the presence of SGI1 integrated in the chromosome suggesting that SGI1 does not provide regulatory factors stronger than LexA-mediated SOS regulation for the transcriptional expression of its master activators SgaDC (Fig. S1). To further confirm the SOS regulation of P*sgaDC*, we performed electrophoretic mobility shift assays with the P*sgaDC* region containing the wild-type and mutated putative LexA binding box (Fig. 2) with increasing concentrations of purified LexA proteins from *Salmonella* or *E. coli* (33). Mobility of the P*sgaDC* probe was delayed by the addition of 1.34 - 2.68 μM LexA from *Salmonella* or *E. coli* (Fig. 3C and Fig. S2B). Nucleotide Substitutions of the essential CTG motif in the P*sgaDC* putative LexA binding box (Fig. 2) completely abolished the binding of LexA (Fig. 3C and Fig. S2B). All these results suggested that the expression of the SGI1 master activator SgaDC is repressed by LexA and activated by the SOS response in both *Salmonella* and *E. coli*. Conversely, the toxin-antitoxin *sgiAT* promoter exhibited a pretty strong activity in the absence of treatment and showed no increase of activity upon SOS induction in both *Salmonella* strains (Fig. S1). A ∼1.5-fold increase of P*sigAT* activity was observed in an *E. coli* background (chemical treatments or SOS^ON^ vs WT) (Fig. S1). Interestingly, the presence of SGI1 in the chromosome seemed to increase P*sgiAT* activity in *S.* Kentucky ST198 and in *E. coli* SOS^ON^ (Fig. S1). The later observations need to be further studied but are in accordance with important self-regulation of toxin-antitoxin systems (34). Confirming our above results and the low weight and associated *P*-value given by the regulatory sequence analysis tools of the P*sgiAT* putative LexA binding site, no mobility shift was detected for the P*sgiAT* probe with purified LexA proteins from *Salmonella* and *E. coli* (Fig. S2C and S2D).

**FIG 3.**
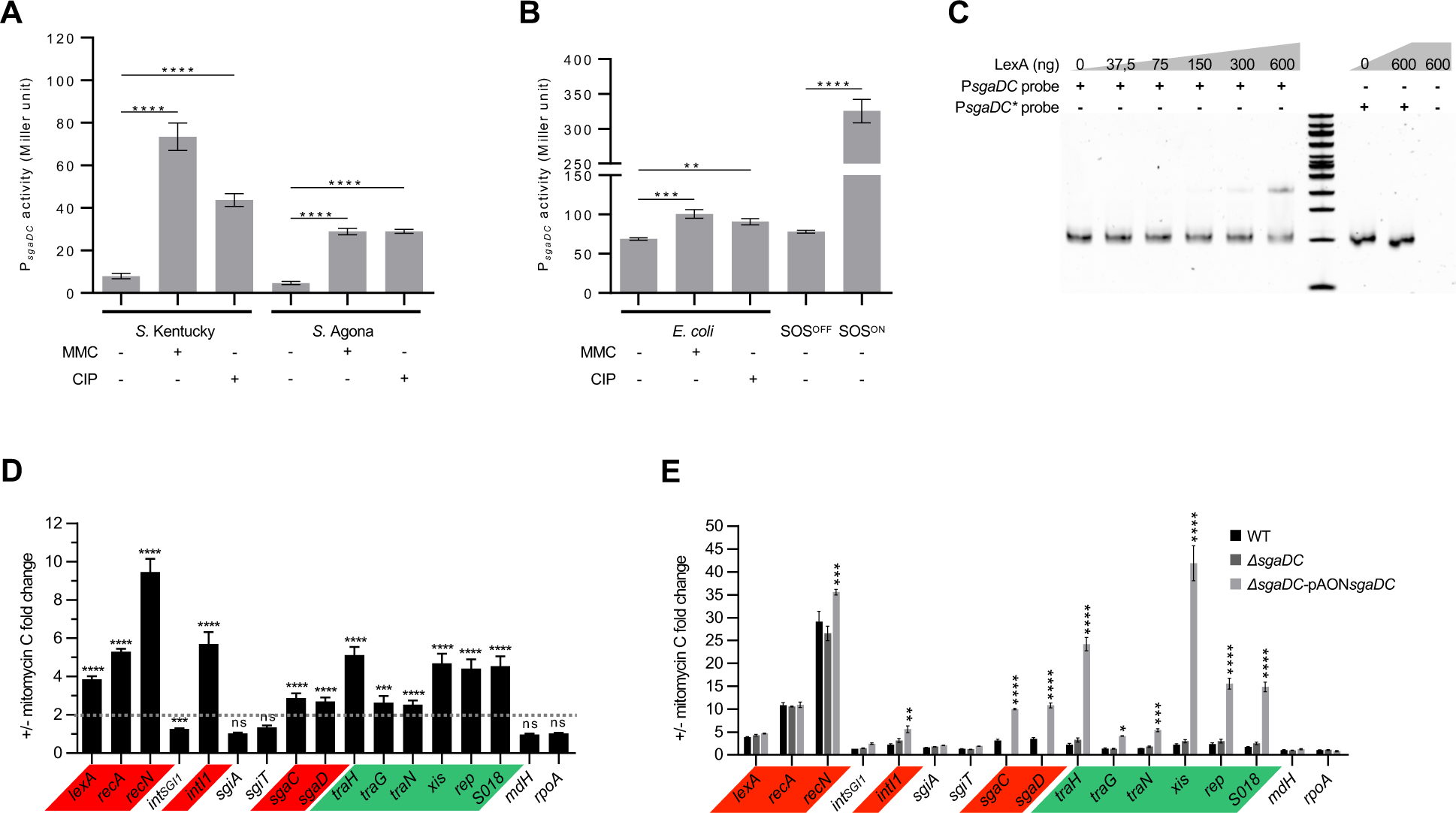
The SOS response controls the expression of the master regulator *sgaDC* and as a consequence the expression of SGI1 essential genes for transfer. (**A**) and (**B**), Promoter activity of P*sgaDC* measured by β-galactosidase activity tests in *S.* Kentucky ST198 strain 11-0799 and *S.* Agona strain 959SA97ΔSGI1 with or without mitomycin C (MMC) or ciprofloxacin (CIP) (**A**) and in *E. coli* WT strain MG1655 with and without MMC or CIP and in MG1655 mutants *lexA3* (SOS^OFF^) and *lexA51* (SOS^ON^) (**B**). The bars represent the mean and standard error of the mean obtained from at least 3 independent experiments, each one assorted of technical duplicates. For each condition, basal β-galactosidase activities were determined using the pQF50 reporter vectors without cloned promoter (data not shown, see source data). One-way ANOVA with Dunnett’s multiple comparison test was performed between induced condition and non-induced. Statistical significance is indicated as followed: ****, *P*<0.0001; **, *P*<0.01. (**C**), Electrophoretic mobility shift assay of the *sgaDC* promoter fragment with increasing quantity (ng) of the *Salmonella* LexA protein from *S.* Agona strain 47SA97. P*sgaDC* and P*sgaDC\** probes contain the native and mutated LexA binding sites, respectively. (**D**), RT-qPCR quantification of gene expression of main SGI1 genes in *Salmonella* (*S.* Agona 47SA97) after treatment or not with mitomycin C. Results are indicated as fold change upon mitomycin C treatment. Two-fold increase in gene expression is indicated with the dotted bar. The bars represent the mean and standard error of the mean obtained from 3 independent experiments, each one assorted of technical triplicates. Genes under the control of LexA and AcaDC/SgaDC are highlighted in red and green, respectively. Statistical significance was determined using multiple *t*-tests with the Holm-Sidak method from the 2^-ΔCT^ values of each mRNA gene level with or without mitomycin C (Fig. S3). Statistical significance is indicated as followed: ****, *P*<0.0001; ***, *P*<0.001; ns, not significant. (**E**), RT-qPCR quantification of gene expression of main SGI1 genes in *E. coli* after treatment or not with mitomycin C. Results are indicated as fold change upon mitomycin treatment. Fold change of gene expression with or without mitomycin C is represented for *E. coli* MG1655::SGI1 (WT), MG1655::SGI1Δ*sgaDC* (Δ*sgaDC*), and MG1655::SGI1Δ*sgaDC* trans-complemented with pAON-*sgaDC* (Δ*sgaDC-* pAON*sgaDC*). Genes under the control of LexA repressors and AcaDC/SgaDC activators are highlighted in red and green, respectively. The bars represent the mean and standard error of the mean obtained from 3 independent experiments, each one assorted with technical duplicates (Fig. S3). Statistical significance was determined using Two-way ANOVA with Dunnett’s multiple comparisons test between Δ*sgaDC* or Δ*sgaDC-*pAON-*sgaDC* compared to WT gene fold-changes. Statistical significance is indicated as followed: **** *P*<0.0001; *** *P*<0.001; ** *P*<0.01; * *P*<0.5; ns not significant.

### All SGI1 genes of the AcaDC/SgaDC regulon are expressed under SOS induction of *sgaDC* expression

To estimate the impact of SOS induction on SGI1 gene expression, we determined the relative expression of the master activator *sgaDC* genes and its downstream regulon in *S.* Agona 47SA97 (carrying SGI1-C, Table S1) using RT-qPCR with or without mitomycin C treatment. As expected, mitomycin C treatment resulted in the induction of the expression of *sgaDC* (2.8-fold), of the SOS regulon genes *lexA, recA* and *recN* as well as the integrase gene *intI1* of the class 1 integron carried by SGI1 (Fig. 2), these later ones all known to be regulated by the SOS response and used here as positive controls (Fig. 3D, genes highlighted in red) (28, 33, 35). Furthermore, mitomycin C treatment resulted in the concomitant expression increase of the AcaDC/SgaDC regulon in SGI1 (*traGHN*, *xis*, *rep*, S018) ranging from 2.5- to 5.1-fold (Fig. 3D, genes highlighted in green). Whilst the toxin-antitoxin genes *sgiAT* showed the higher level of expression among SGI1-tested genes in *Salmonella* (Fig. S3A), we confirmed that their expression is not regulated by the SOS response in *Salmonella* (Fig. 3D). In addition, the SGI1 integrase gene *int*_SGI1_ remained constitutively expressed without any change under mitomycin C treatment. Gene expression was also assessed in the *E. coli* background MG1655::SGI1 (carrying SGI1-C from 47SA97; Table S1), as well as derivatives MG1655::SGI1Δ*sgaDC* with or without *trans*-complementation with pAON*sgaDC* vector (low copy plasmid containing *sgaDC* under the control of its native promoter; Table S2) (Fig. 3E). In the *E. coli* background, mitomycin C treatment in the artificial context of multi-copies of *sgaDC* (SGI1Δ*sgaDC*-pAON*sgaDC*) resulted in a 10-fold induction of *sgaDC* expression and a strong up-regulation of all genes of the AcaDC/SgaDC regulon in SGI1 (Fig. 3E, Fig. S3). Altogether, these results showed that induction of the SOS response results in expression activation of SGI1 master activator genes *sgaDC* that are actually repressed by the SOS master regulator LexA.

The excision and replication of SGI1 to ∼6-12 extrachromosomal circular copies have been previously shown to be activated by SgaDC in the presence of helper IncC plasmid and to be actually responsible of the destabilization of the helper IncC plasmid (17–19). Although IncC plasmids were absent in these SOS induction experiments by mitomycin C, we decided to quantify SGI1 excision and determine its copy number using real-time quantitative PCR. Induction of the SOS response in *E. coli* strain MG1655::SGI1 did not allow to detect the excision or replication of SGI1 (Table 2). In the artificial context of *sgaDC* overexpression (SGI1Δ*sgaDC*-pAON*sgaDC*), although the SGI1 *xis* (excisionase) and *rep* genes were overexpressed 41- and 15-fold, respectively, SGI1 excision remained at a low level, occurring in 6% (empty *attB* site per chromosome) with mitomycin C treatment, and SGI1 did not seem to replicate (copy number remained at 1 copy per chromosome) (Table 2). SGI1 remaining at 1 copy per chromosome confirmed that SOS induction of expression of *sgaDC* and its SGI1 regulon is not due to increase of the SGI1 copy number in these conditions (Fig. 3E and Table 2). These results are in agreement with previously published ones (17–19).

**TABLE 2.** SGI1 copy number and excision of SGI1 in *E. coli* MG1655::SGI1 (WT), MG1655::SGI1Δ*sgaDC* (Δ*sgaDC*), and MG1655::SGI1Δ*sgaDC* trans-complemented with pAON-*sgaDC* (Δ*sgaDC-*pAON*sgaDC*) with or without mitomycin C induction

| SGI1 | MMC | SGI1 copy<br>number<sup>a</sup><br>(*xis*/chr) | Excised SGI1<sup>a</sup><br>(*attP*/chr) | Empty *attB*<sup>a</sup><br>(*attB*/chr) | +/- MMC |  |
| --- | --- | --- | --- | --- | --- | --- |
|  |  |  |  |  | fold change<br>of gene expression<sup>b</sup> |  |
|  |  |  |  |  | *xis* | *rep* |
| WT | - | 1.05 ± 0.04 | ND | ND | NA | NA |
| WT | + | 1.07 ± 0.03 | ND | ND | 2.2 ± 0.2 | 2.4 ± 0.3 |
| SGI1Δ*sgaDC*-pAON*sgaDC* | - | 1.06 ± 0.04 | 0.0013 ± 6.2 10<sup>-5</sup> | 0.0002 ± 8.9 10<sup>-6</sup> | NA | NA |
| SGI1Δ*sgaDC*-pAON*sgaDC* | + | 1.06 ± 0.03 | 0.13 ± 0.004 | 0.06 ± 0.002 | 41.9 ± 3.8 | 15.6 ± 1.2 |

<sup>a</sup>The values represent the mean +/- the standard error of the mean obtained from 3 independent experiments, each one assorted of a technical triplicate. ND and NA stand for non-detected and non-applicable, respectively. SGI1 copy number, excised SGI1 and empty chromosomal attachment site (*attB*) correspond to the mean of ratio between the targets, *xis* gene, *attP*, *attB*, respectively, and 2 distinct chromosomal genes (*mdh* and *rpoA*) as detailed in methods section.
<sup>b</sup>Fold changes of essential genes expression for excision (*xis*) and replication (*rep*) of SGI1 have been determined on mRNA extractions performed at the same time of total DNA extractions from the same 3 independent cultures (see also in Fig. 3E).

### SOS induction upon IncC plasmid acquisition promotes an early transfer of SGI1

The incompatibility phenotype between IncC plasmid and SGI1 is strongly related to SGI1 replication and requires to maintain antibiotic selection pressure on both elements before SGI1 mating assays to ensure the largest SGI1^+^/IncC^+^ donor populations (17, 18, 20). Here, we developed a novel approach based on tripartite conjugation assay to study the impact of IncC plasmid conjugative entry in SGI1 donor cells. Briefly, an *E. coli* strain MG1655 carrying the conjugative helper plasmid (IncC^+^, donor #1) was mixed with a second *E. coli* strain harbouring SGI1 (MG1655::SGI1, donor #2) and a third *E. coli* strain, the rifampicin-resistant (Rif^R^) J5-3 recipient (see Methods). The conjugation frequencies of the IncC plasmid and SGI1 were measured at various time points after initial contact between donors and recipient. In each experiment, IncC plasmid transfer were similar towards SGI1^+^ donor or empty Rif^R^ recipient strains (Table S4; compare R16a transconjugants and SGI1^+^/R16a^+^ donors). In a WT MG1655 *E. coli* background, IncC transfer frequency quickly reached ∼10^-3^ (after 45 min contact) and remained stable along the experiment (Fig. 4A). SGI1 transfer frequency was below the detection limit (10^-7^) until a burst of SGI1 transfer (1.9 10^-2^ ± 5.2 10^-3^; after 1 h 45 min mating) (Fig. 4A). At 4h mating, SGI1 transfer frequency has increased to ∼10^0^ (Fig. 4A). The same results were obtained with *E. coli* MG1655::SGI1 *lexA3* (SGI1-C from 47SA97, SOS^OFF^) as donor #2 (Table S4) that are in agreement with β-galactosidase reporter assays using SOS^OFF^ *E. coli* strain (Fig. 3B). Using *E. coli* MG1655::SGI1 *lexA51* (SGI1, SOS^ON^) as donor #2, whilst the temporal curve of IncC transfer frequency was comparable to that in Fig. 3d, it is striking that the burst of SGI1 transfer occurred 30 min earlier compared to its transfer from *E. coli* WT background MG1655::SGI1 and reached more rapidly a plateau at ∼10^-1^ (Fig. 4B). This resulted in a 4.5-fold increase of SGI1 transfer at 1 h 45 min in the SOS^ON^ context compared to its transfer from *E. coli* WT background MG1655::SGI1 (Fig. 4D). On the other hand, when using *E. coli* MG1655::SGI1Δ*sgaDC* as donor #2 in our tripartite conjugation assay (Fig. 4C), we observed a higher transfer frequency of IncC plasmid and a lower SGI1 transfer frequency at 1 h 45 min compared to *E. coli* WT background MG1655::SGI1 (Fig. 4D). These results clearly indicated that permanent SOS induction (SOS^ON^) induces early transfer of SGI1 following IncC plasmid acquisition by MG1655::SGI1 donor cells. Altogether, these results confirmed the importance of *sgaDC* for SGI1 transfer and that deletion of *sgaDC* abolishes the incompatibility with IncC plasmid (17, 18, 22).

**FIG 4.**
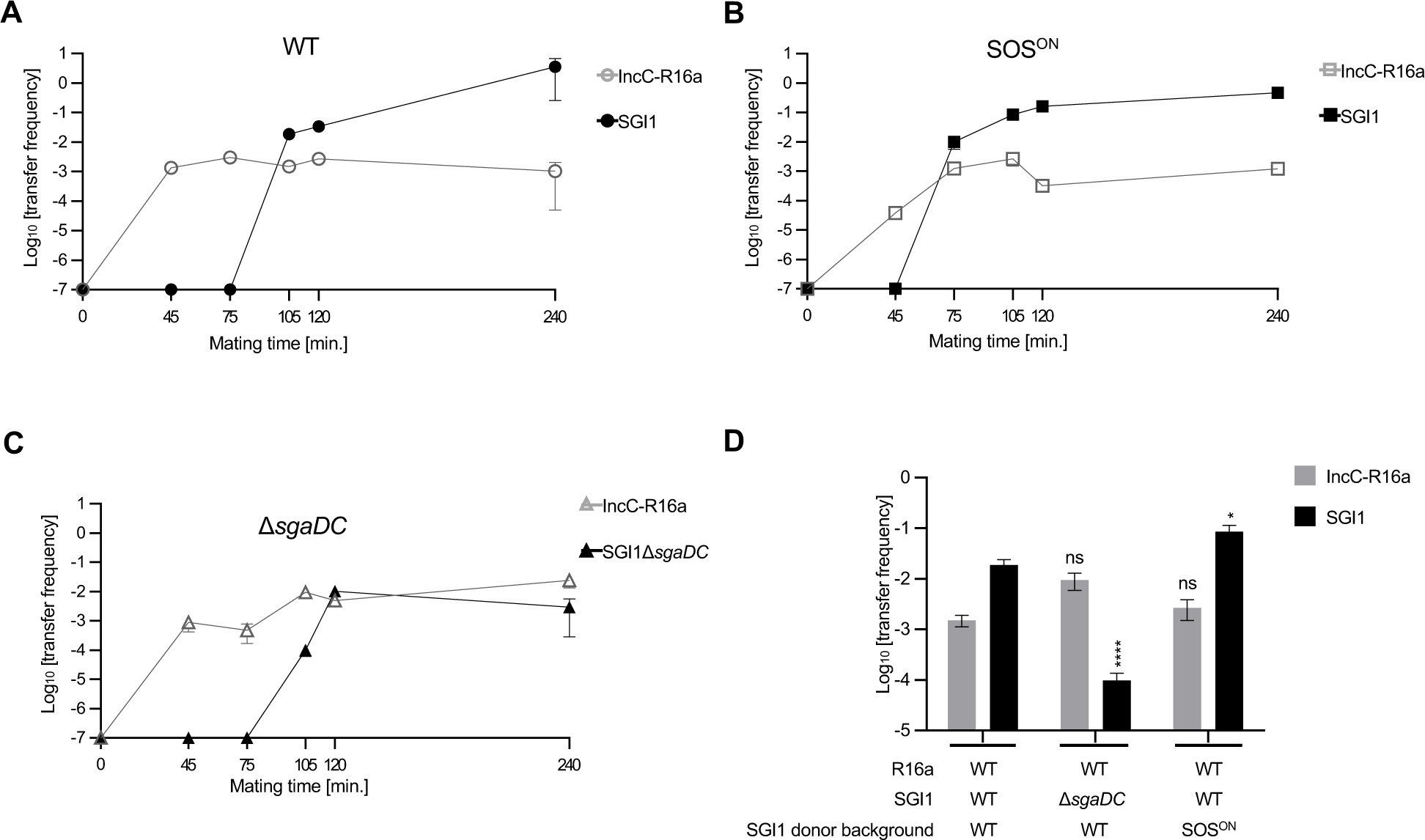
Time-dependent conjugative transfer of SGI1 and IncC plasmid in tri-partite conjugation assays. IncC-R16a and SGI1 transfer frequencies (empty and filed symbols, respectively) as a function of mating time in tripartite conjugation assays. In all experiments, IncC plasmid R16a and SGI1 were carried by two distinct *E. coli* MG1655 donor strains (R16a in donor #1 and SGI1 or derivatives in donors #2). Donors #1 and #2 were mixed together with a third *E. coli* recipient strain, the rifampicin-resistant J5-3 in a 1:1:1 ratio. In all experiments, the IncC plasmid R16a was carried by WT *E. coli* strain MG1655 (donor #1). (**A**), donor #2: WT *E. coli* strain MG1655::SGI1. (**B**), donor #2: *E. coli* MG1655::SGI1 *lexA51* mutant (SGI1, SOS^ON^). (**C**), donor #2: WT *E. coli* strain MG1655::SGI1Δ*sgaDC*. (**D**), IncC-R16a and SGI1 Transfer frequencies after 1 h 45 min of mating in tripartite conjugation assays from panels (A), (B), and (C). The bars represent the mean and standard error of the mean obtained from at least 3 independent experiments. Statistical significance was determined using One-way ANOVA with Dunnett’s multiple comparisons test on the logarithm of the values between WT *E. coli* strain MG1655::SGI1Δ*sgaDC* (C) or MG1655::SGI1 *lexA51* (B) compared to WT *E. coli* strain MG1655::SGI1 (A). Statistical significance is indicated as followed: ****, *P*<0.0001; *, *P*<0.5; ns not significant.

## DISCUSSION

The recent characterization of the SgaDC regulon revealed the same binding pattern as AcaDC both on SGI1 and on the IncC plasmid resulting in the expression of the transfer genes (17). While AcaDC replacement by SgaDC appears fully functional for SGI1 and IncC plasmid transfers, SgaDC is essential for SGI1 replication and subsequently the destabilization of the IncC plasmid (17, 18, 22). Altogether, these recent findings and our study argued for an earlier crucial role of SgaDC in the complex regulatory interaction between SGI1 and IncC plasmid. We propose a model of early temporal regulation of SGI1 gene expression under the control of SgaDC due to the transient SOS induction following the conjugative entry of IncC plasmid in SGI1 bearing cells (Fig. 5). As shown in this study, we demonstrated that the conjugative entry of IncC plasmid activates the SOS response in recipient cells (Fig. 1) thus suppressing the repression of LexA on *sgaDC* in the recipient SGI1 cells (Fig. 3). This results in the expression of *sgaDC* and its SGI1-encoded regulon (*xis*, *rep*, *traNGH* and S018) (Fig. 3D and 3E). Moreover, in the absence of IncC plasmid, overexpression of *sgaDC,* although upregulating its SGI1 regulon, did not result in strong SGI1 excision nor replication (Fig. 3E and Table 2), confirming that Inc-encoded factors, *e.g.* AcaDC and probably others are needed (17). Thanks to the novel tripartite conjugation approach, we confirmed the role of the SOS induction in the early timing of SGI1 transfer following the conjugative entry of IncC plasmid (Fig. 4).

**FIG 5.**
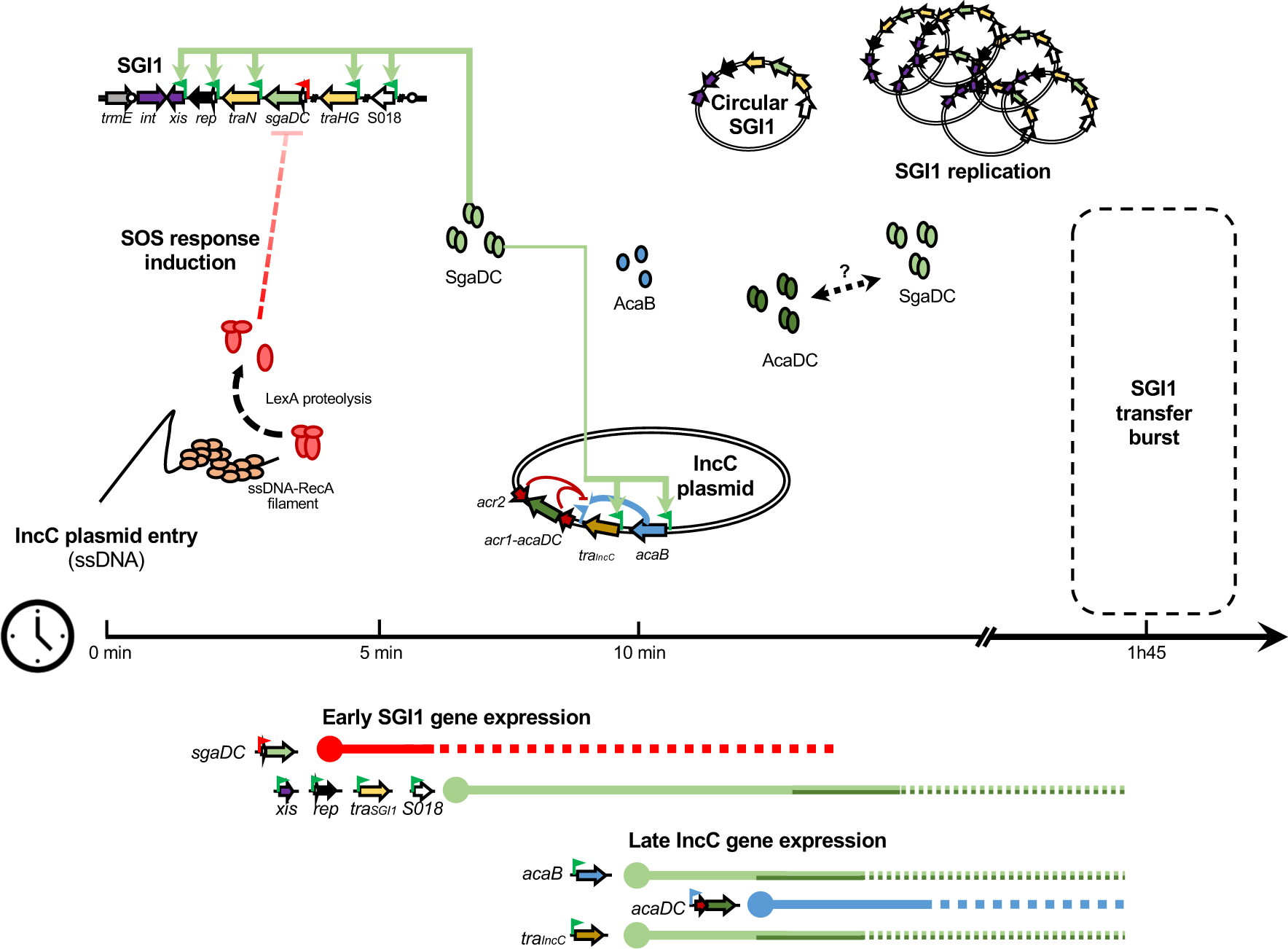
Proposed model of early timing of SGI1 gene expression for transfer upon IncC plasmid entry. Genes of interest are represented by arrows in SGI1 (located in the chromosome) and IncC plasmid. Gene and protein functions are color-coded as follow: yellow/brown, transfer; blue, light and dark green, transcriptional activators; red, repressors; and purple, recombination. Promoters are represented by color-coded flags according to the corresponding repressor/activator: red, LexA repressed; green, AcaCD/SgaCD-activatable; and blue, AcaB activatable. The red blocked dashed arrows represented the removal of the inhibition of the *sgaDC* promoter by the LexA repressor. The light green arrows represent the activation of gene expression mediated by SgaDC. A bidirectional black dashed arrow indicates potential interactions between AcaDC and SgaDC. The proposed model of temporal gene regulation depicted at the bottom of the figure is based on all the results obtained in this study and other findings on the complex SGI1-IncC biology, recently published^10,24^. Gene expression is shown as color-coded solid/dotted lines according to the corresponding activators along the time scale (see text for timing explanation).

The proposed timing in Fig. 5 is based on: (i) conjugative DNA transfer rate of ∼45 kb/min suggesting transfer of the ∼160 kb IncC plasmid should be achieved in ∼3 min, (ii) plasmidic-ssDNA has been shown to have a lifetime of ∼5 min, before (iii) conversion to double strand DNA (dsDNA) circular form that arises between 5 to 10 min after conjugative entry in the recipient cells (36). After initiation of transfer, IncC plasmid should be in a dsDNA circular form within ∼10-12 min. Thus, the transient SOS induction could occur during ∼5 min after initiation of IncC entry, resulting in early expression of *sgaDC* and of its SGI1 regulon (*traGHN*, *xis*, *rep*, S018) probably before IncC conversion to dsDNA and IncC gene expression. Soon after IncC dsDNA conversion, SgaDC could activate expression of *acaB* and the SgaDC/AcaDC regulon (e.g. *tra*_IncC_) of IncC plasmids. Hence, expression of *acaDC* may happen later on *via* AcaB activation, as recently published (17). The low-level expression of SgaDC upon SOS induction (Fig. 3D) coupled to the presence of an IncC plasmid to achieve high level of SGI1 excision and replication (Table 2) could represent the two mandatory signals for SGI1 transfer. Thus, in the case of SOS response induction independently of IncC conjugative entry (e.g. ciprofloxacin), SGI1 probably remains integrated in the chromosome to ensure its maintenance.

Our study shed light on the role of the SGI1 master activator *sgaDC* acting as the sensor for IncC plasmid entry through the SOS regulation of its promoter as previously suggested (17). This regulation complies with the crucial role of the SGI1 master activator *sgaDC* for SGI1 excision, replication, conjugative mobilisation and incompatibility with plasmids of the IncC family(17–19). Besides their expression, several biological interactions between SgaDC and AcaDC master activators remain to be elucidated (mRNA half-life, translation, complex stability, potential chimeric complexes) that can play a role in the intimate cross-talk between SGI1 and its helper IncC plasmids. SOS induction during bacterial conjugation may likely impact a wide range of recipient genomes, thus promoting the dissemination of the SGI1 or related islands with the consecutive spread of antibiotic resistance genes in diverse bacterial pathogens (26, 28, 33, 35). The conjugative transfer as well as the SOS response may therefore constitute suitable targets for co-treatment with antibiotics in order to prevent dissemination and exchange of antibiotic resistance genes (26, 28, 37).

## MATERIALS AND METHODS

### Bacterial strains and growth conditions

A list of strains used in this study is provided in Table S1. *Salmonella* and *E. coli* strains were routinely grown in lysogeny broth (LB) at 37°C (or 30°C for thermosensitive vectors) under agitation at 180 rpm. Cultures on solid media were realized using LB or *Salmonella*-*Shigella* (SS) media. Antibiotics were used at the following concentrations: ampicillin (100 µg/mL, Amp), chloramphenicol (30 µg/mL, Chl), kanamycin (50 µg/mL, Kan), streptomycin (50 µg/mL, Str), and tetracycline (10 µg/mL, Tet), and rifampicin (250 µg/mL, Rif).

### Plasmids, primers and bacterial construction

All plasmids and primers used in this study are listed in Tables S2 and S3, respectively. To quantify promoter activities by β-galactosidase assays, different promoter regions were amplified using the Phusion® high-fidelity DNA polymerase (NEB) from purified genomic DNA. PCR products digested by *Sph*I and *Hin*dIII were cloned into pQF50-chl and/or pQF50-Amp plasmids using T4 DNA ligase (NEB) to generate the different pMP plasmids (38, 39). pMPs plasmids were transformed into *S.* Agona strains 959SA97ΔSGI1, 47SA97 harbouring SGI1-C, and the following strains harbouring or not SGI1-C (originating from 47SA97): *S.* Kentucky ST198 strain 11-0799, *E. coli* strains MG1655, MG1655 *lexA3*, and MG1655 *lexA51*. The *lexA* ORF of *S.* Agona strain 47SA97 was amplified using primers EMSA-LexAS-p15b_F and EMSA-LexAS-p15b_R. The PCR product was digested by *Nde*I and *Xho*I, ligated into *Nde*I/*Xho*I-digested pET15b expression vector and electroporated into competent *E. coli* strain BL21(DE3)-pLysS. SGI1-CΔ*sgaCD* was initially constructed in *S.* Agona strain 47SA97 using the one-step chromosomal gene inactivation method using primers Rec-delsgaCD_F and Rec-delsgaCD_R primers (40). SGI1-C and SGI1-CΔ*sgaCD* were further transferred in different strains by a two-step conjugation protocol. Briefly, the IncC plasmids R55 or R16a was first introduced by conjugation in the SGI1 host strain. In a second conjugation, SGI1 or derivatives were transferred to the required strain.

### Tripartite conjugation assays

Conjugation experiments were carried out on filters using simultaneously two distinct donor *E. coli* strains, the first (MG1655; donor #1) harbouring the IncC plasmid R16a, the second one (MG1655::SGI1; donor #2) carrying SGI1 and a third recipient *E. coli* strain (J5-3 rifampicin-resistant). Briefly, the IncC plasmid R16a, SGI1-C or SGI1-CΔ*sgaDC* were previously introduced by conjugation in the different donor *E. coli* MG1655 derivatives (WT, *lexA3*, and *lexA51*). Following overnight cultures of donor and recipient strains at 37°C with the appropriate antibiotics, cultures were refreshed 1:100 in 10 mL LB medium without antibiotics and grown up to OD_600_ ∼ 0.8 at 37°C under agitation. Donor and recipient cells were mixed in a 1:1:1 ratio in 3 mL final volume, and concentrated in 200 µL by smooth centrifugation for 3 min at 765 g. Mating mix were applied on 0.045µm mating filter on 37°C-prewarmed LB plates and then incubated at 37°C. At different time points of contact, up to 4h contact, filters were resuspended in 1 mL LB medium, and 10-fold serial dilutions were plated on LB media with the appropriated antibiotics to determine SGI1 and R16a transconjugants, SGI1 donors carrying R16a and SGI1, and recipients. Transfer frequencies correspond to the ratio of transconjugants/donors.

### β-galactosidase tests in recipient population

The specific activation of the SOS response following plasmid entry in recipient cells was determined for conjugative plasmids Rsa (IncW, positive control), R55 (IncC) and RA1 (IncA) using the protocol previously described by Baharoglu et al. with minor adaptations. *E. coli* strain TOP10 (*recA^-^*and Δ*lacZ*; Table S1) harbouring one of the conjugative plasmids above and *E. coli* strain MG1655 carrying reporter vectors pMP002 or pMP010 (*lacZ* expression under the promoter P*recN*) were used as donor and recipient strain, respectively (26). The protocol of filter mating described above was applied with a 1:1 ratio of donor and recipient cells. Ten-fold serial dilutions of mating mixes were plated on LB media with the appropriated antibiotics to determine transconjugants, donors, and recipients. Transfer frequencies correspond to the ratio of transconjugants/donors.

β-galactosidase activity tests were performed as described previously (see also section below) (26, 41). Briefly, the basal β-galactosidase activity per recipient cell was determined in mating assays with an empty donor strain (without conjugative plasmid) at different time points (1h and 2h; in absence of SOS response induction by ssDNA entry).

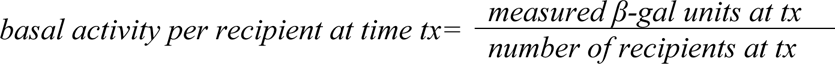

For each plasmid, the β-galactosidase activity tests were performed at transfer frequencies ranging from 10^-4^-10^-3^ (1h or 2h filter mating). To determine the specific β-galactosidase activity per transconjugant at a given time point, the basal activity of the recipient population (without plasmid) was removed from the β-galactosidase activity in mating assay with a conjugative plasmid to assign the remaining activity to the transconjugant population.

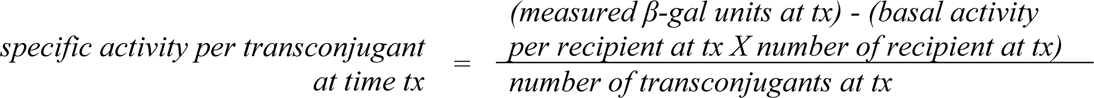

Finally, the results were represented as SOS induction ratios for each plasmid corresponding to

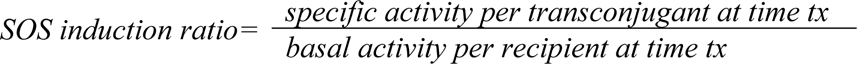

### β-galactosidase assays

The quantification of β-galactosidase activity in Miller units was realised as described by Miller adapted to 96 well plates as follow (41). For each sample, 2 dilutions of bacterial lysates were used in technical triplicate. To assess the induction of the SOS response during conjugation, 500 µL of the mating assay was lysed and used to quantify the β-galactosidase activity at 12.5- and 25-fold final dilutions. To quantify promoter activities, the pMP-carrying strains were grown with or without mitomycin C (200 ng/mL) or ciprofloxacin (used at concentration 100-fold below MICs) up to OD_600_ ∼ 1.5. One mL of bacterial culture was centrifuged and used to quantify the β-galactosidase activity at 40- and 100-fold final dilution.

### Electrophoretic mobility shift assay (EMSA)

Overproduction of LexA proteins of *E. coli* strain MG1655 or *S.* Agona were carried out in *E. coli* strain BL21(DE3)-pLysS using pUA1170 or pET15b-*lexA_Salmonella_*, respectively. LexA production was induced with 5mM IPTG in 200 mL LB medium at OD_600_ ∼ 0.5 during 4 hours at 37°C under shaking at 250 rpm. The purification was performed using TALON® Metal Affinity Resin from (Clontech Laboratories, Inc, 635501) with HisTALON™ Buffer Set (Clontech Laboratories, 635651) following manufacturer’s instructions. The elution fractions containing LexA proteins were pulled, dialyzed with buffer (20mM Tris-HCl pH8, 50mM KCl, 1mM EDTA) and concentrated with Vivaspin Turbo 15 RC, 10,000 MWCO (Sartorius, VS15TR01). The DNA probes P*_sgaCD_* and P*_sgiAT_* were amplified on total DNA of *S.* Agona strain 47SA97 using primers EMSA-sgaCDbox_F/EMSA-sgaCDbox_R and EMSA-sgiATbox_F/EMSA-sgiATbox_R, respectively (Table S3). Nucleotide changes from CTG to AGT in the potential LexA binding boxes to produce mutated probes were performed by overlap PCR using primers above and primers EMSA-mut-sgaCDbox_F/EMSA-mut-sgaCDbox_R and EMSA-mut-sgiATbox_F/EMSA-mut-sgiATbox_R for DNA probes P*_sgaCD_\** and P*_sgiAT_\**, respectively (Table S3). EMSAs were realized using the Electrophoretic Mobility-Shift Assay Kit (Thermo Fisher Scientific, E33075). Briefly, 40 ng of DNA probes were mixed with different amounts of LexA proteins ranging from 0 to 600 ng in binding buffer E in a final volume of 10 µl and incubated for 20 min on ice. Samples were separated in 6% non-denaturing Tris-glycine polyacrylamide gels and visualised with SYBR® Green following the manufacturer’s protocol.

### Quantification of gene expression by RT-qPCR

Bacterial strains were grown in triplicate cultures with or without mitomycin C at 200 ng/mL in 10 mL LB medium at 37°C under agitation until reaching OD_600_ ∼ 1. RNA extractions were performed from 1 ml LB culture using the Direct-zol RNA MiniPrep Plus (Zymo Research, ZR2073). Total RNAs were reverse-transcribed into cDNAs using iScript™ cDNA Synthesis Kit (BioRad, 1708891). The qPCR assays were performed with a Biomark™ HD system (Fluidigm) and primers listed in Table S3. According to the manufacturer’s instructions, cDNA samples were diluted 10-fold and pre-amplified with a mix of all primers using the Pre-Amp master Mix (Fluidigm, 100-5581) and amplification conditions: 2 min at 95 °C, followed by 14 cycles of 15 s at 95 °C and 4 min at 60 °C, and a final hold at 4 °C. Samples were treated with exonuclease I to removed primers (NEB, M0293L) and diluted 20-fold in Tris-EDTA. Pre-amplified cDNA samples and primer pairs were loaded on the Fluidigm 96.96 or 48.48 Dynamic Array™ IFC and run in the Biomark™ HD system using the following program: 70 °C for 40 min, 60 °C for 30 s, 95°C for 60 s, 30 cycles of 96 °C for 5 s, 60 °C for 20 s, followed by a melting curve. Data were analysed with Fluidigm Real-Time software to determine the cycle threshold values (Ct). For each sample, gene expression was normalized to the geometric mean of the housekeeping genes *rpoA* and *mdH* to obtain ΔCt value. Both of these housekeeping genes have previously been used in SOS induction studies and their expression is thus known to be unaffected (27, 42). Fold changes of gene expression (2^-ΔΔCT^) correspond to the ratio of the 2^-ΔCT^ means with and without mitomycin C treatment.

### Determination of the SGI1 copy number and excision rate by qPCR

Total DNA extractions were realized simultaneously to RNA extractions for *E. coli* strains MG1655::SGI1, MG1655::SGI1-CΔ*sgaDC*, and MG1655::SGI1Δ*sgaDC*/pAON*sgaDC* using the NucleoSpin Tissue Mini kit (Macherey Nagel, 740952.50). Primers used to quantify the chromosome and different forms of SGI1 are listed in Table S3. qPCR was carried out with the Bio-Rad® CFX96 Touch™ with the iQ™ SYBR® Green Supermix (Bio-Rad, #1708882) with the following amplification conditions: 5 min at 95 °C, 42 cycles of 15 s at 95 °C and 1 min at 60 °C followed by a melting curve. SGI1 copy number, percentage of excised SGI1 (*attP*) and empty chromosomal attachment site (*attB*) correspond to the ratio between the 2^-ΔCt^ means of targets, *xis* gene, *attP*, *attB*, respectively, and the 2^-ΔCt^ means of 2 housekeeping chromosomal genes (*mdh* and *rpoA*) for each DNA sample.

### Statistical analysis

Statistical analyses were performed with Prism 6.0 software (GraphPad). Analyses were performed on data from three to six independent experiments depending on the experiments. The different statistical tests used are indicated in figure legends. Significance is indicated by *P*-values: ns, non-significant; ∗, *P*<0.05; ∗∗, *P*<0.01, ∗∗∗, *P*<0.001; and ∗∗∗∗, *P*<0.0001.

### Data availability

Raw read data and draft genome assemblies of *Salmonella* strains (11-0799 and 47SA97) used in this study have been deposited in the European Nucleotide Archive under bio-project PRJEB52018. Sanger sequencing data confirming all plasmid constructions and mutants as well as experimental source data are available from the authors upon reasonable request.

## ACKNOWLEDGEMENTS

The authors would like to acknowledge Pr. Laurence Van Melderen (ULB, Brussels, Belgium) for providing *E. coli* strains, Dr. Pierre Germon (INRAE, Nouzilly, France) for expert technical assistance in Fluidigm experiments and analysis, Maxime Branger for genome assemblies and annotation, and Dr. Olivier Grépinet (INRAE, Nouzilly, France) for technical assistance in β-galactosidase reporter assays. This work was supported by the One Health European Joint Program (Grant number 773830) under the Fullforce project, and by public funds from the French National Research Institute for Agriculture, Food, and Environment (INRAE), and from Institut National de la Santé et de la Recherche Médicale (Inserm).

MCP is supported by a PhD fellowship from the microbiology and food chain division of INRAE and the French administrative region Centre-Val-de-Loire.

## Author contributions

M.C.P., A.C., and B.D. conceived the study. M.C.P. performed all experiments and analysis with the assistance of K.P. and S.D.R. A.C., and B.D. supervised aspects of the study and provided essential expert analysis and funding. B.D. wrote the original draft of the manuscript with input from M.C.P. and corrections from all authors. All authors contributed to the interpretation of the results, read and approved the final manuscript.

## Additional information

### SUPPLEMENTAL MATERIAL

**FIG S1** Level of P*sgaDC*, P*sgiAT*, P*acaDC* and P*recN* activity in the presence or absence of SGI1 in *Salmonella* and *E. coli* with or without mitomycin C (MMC) or ciprofloxacin (CIP). Promoter activity of P*sgaDC* (**A**) and (**B**), P*sgiAT* (**C**) and (**D**), P*acaDC* (**E**) and (**F**), and P*recN* (**G**) and (**H**) were assessed in absence 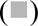 or presence 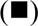 of SGI1 integrated in the chromosome in wild-type *S.* Kentucky ST198 strain 11-0799, *S.* Agona strains 959SA97ΔSGI1 and 47SA97 SGI1-C (A), (C), (E), (G) and *E. coli* strains MG1655 and derivatives *lexA3* (SOS^OFF^) and *lexA51* (SOS^ON^) (B), (D), (F), (H) with or without mitomycin C (MMC) or ciprofloxacin (CIP) to induce SOS response as well as in *E. coli* mutants *lexA3* (SOS^OFF^) and *lexA51* (SOS^ON^) (B), (D), (F), (H). The bars represent the mean and standard error of the mean obtained from at least 3 independent experiments, each one assorted of with technical duplicates. Two-way ANOVA with Sidak’s multiple comparisons test was performed to compare the presence/absence of SGI1. One-way ANOVA with Dunnett’s multiple comparison test was performed between induced condition and non-induced and only shown if significant except for (A) and (B) (see Figs 3A and 3B, respectively). Statistical significance is indicated as follow: ****, *P*<0.0001; ***, *P*<0.001; **, *P*<0.01; *, *P*<0.05; ns, not significant.

**FIG S2** Electrophoretic mobility shift assay of the P*sgaDC* and P*sgiAT* regions with *Salmonella* or *E. coli* LexA proteins. (**A**), Partial sequences of *sgaDC* and *sgiAT* promoters are indicated showing putative -10, -35 regions, and LexA binding box. The 3 essential nucleotides (CTG) of LexA binding boxes that have been substituted in the mutated probes are indicated by stars in Fig. 2. (**B**), Electrophoretic mobility shift assay of the *sgaDC* promoter fragment from *S.* Agona strain 47SA97 carrying SGI1-C and increasing quantities of the LexA protein from *E. coli* MG1655. P*sgaDC* and P*sgaDC\** probes contain the native and mutated LexA binding sites, respectively. (C) and (D), Electrophoretic mobility shift assay of the *sgiAT* promoter fragment from *S.* Agona strain 47SA97 carrying SGI1-C and increasing quantities of LexA protein purified from (**C**) *S.* Agona strain 47SA97 or (**D**) *E. coli* strain MG1655. P*sgiAT* and P*sgiAT\** probes contain the native and mutated LexA-binding sites, respectively.

**FIG S3** Relative gene expression with or without mitomycin C treatment to induce the SOS response. Box plots of the 2^-ΔCT^ values of mRNA gene level with or without mitomycin C in *S.* Agona strain 47SA97 harboring chromosomally-integrated SGI1 (**A**), in *E. coli* MG1655::SGI1 (**B**)*, E. coli* MG1655::SGI1Δ*sgaDC* (**C**), and in *E. coli* MG1655::SGI1-CΔ*sgaDC* trans-complemented with pAON*sgaDC* (**D**). The derivatives fold changes of gene expression (2^-ΔΔCT^) are shown in Fig. 3D and E. Genes under the control of LexA repressors and AcaDC/SgaDC activators are highlighted in red and green, respectively. Boxes extend from the 25^th^ to 75^th^ percentile of each group’s distribution; horizontal inner lines indicate median values; vertical extending lines denote the minimum and maximum values obtained from 3 biological independent experiments, assorted of technical triplicates for each. Statistical significance was determined using multiple *t*-tests with the Holm-Sidak method from the 2^-ΔCT^ values of mRNA gene level to compare with and without mitomycin C treatment. Statistical significance is indicated as follow: ****, *P*<0.0001; ***, *P*<0.001; **, *P*<0.01; *, *P*<0.05; ns, not significant.

